# Modulation of Aplnr signaling is required during the development and maintenance of the hematopoietic system

**DOI:** 10.1101/2020.06.15.145649

**Authors:** Melany Jackson, Antonella Fidanza, A. Helen Taylor, Stanislav Rybtsov, Richard Axton, Maria Kydonaki, Stephen Meek, Tom Burdon, Alexander Medvinsky, Lesley M. Forrester

**Affiliations:** MRC Centre for Regenerative Medicine, University of Edinburgh, 5 Little France Drive, Edinburgh EH16 4UU; Institute for Stem Cell Research, Centre for Regenerative Medicine, 5 Little France Drive, Edinburgh EH16 4UU; Roslin Institute, University of Edinburgh, Easter Bush, Midlothian, EH25 9RG

## Abstract

Apelin receptor (Aplnr/Agtrl1/Apj) marks a transient cell population during the differentiation of hematopoietic progenitor cells (HPCs) from pluripotent stem cells but the function of this signalling pathway during hematopoietic development both in vitro and in vitro is poorly understood. We generated an Aplnr-null mouse embryonic stem cell (mESC) line and demonstrated that they are significantly impaired in the production of HPCs indicating that the Aplnr pathway is required for their formation. Using Aplnr-tdTomato reporter mESCs we demonstrated that is expressed in a population of differentiating mesodermal cells committed to a hematopoietic and endothelial fate. Activation of this signaling pathway by the addition of the Apelin ligand to differentiating ESCs has no effect on the production of HPCs but the addition to *ex vivo* AGM cultures impaired the generation of long term reconstituting hematopoietic stem cells and appeared to drive myeloid differentiation. Taken together, our data suggest that the Aplnr pathway is required for the generation of cells that give rise to HSCs during development but its subsequent down regulation is required for their maintenance.

**HIGHLIGHTS:**

- Hematopoietic differentiation is impaired in Aplnr-null ESCs
- Aplnr-tdTomato reporter marks a subpopulation of ESC-derived mesoderm.
- Aplnr signaling drives the maturation of lineage-committed myeloid progenitors
- In AGM explant cultures HSC activity is reduced in the presence of Aplnr ligands.

## INTRODUCTION

Definitive hematopoietic stem cells (HSCs) comprise a rare but potent cell population in the adult bone marrow that is capable of repopulating the entire blood system upon transplantation. HSCs arise through a complex process, precisely coordinated at a number of anatomical sites throughout embryonic development. In the mouse embryo the first primitive wave of hematopoietic development originates in the yolk sac around embryonic day (E) 7.25 and gives rise to embryonic erythrocytes, megakaryocytes and macrophages^1^. The second wave, also originating from the yolk sac from E8.25, gives rise to erythromyeloid progenitors (EMP) and are distinguished by their ability to generate granulocytes^2^. The third wave originates in the ventral region of the developing dorsal aorta within the aorta-gonad mesonephros (AGM) at E10.5-E11.5 and generates HSCs that can repopulate the entire hematopoietic system upon transplantation^3, 4^. HSCs arise from a subpopulation of endothelial cells that co-express hematopoietic markers such as CD41 and Kit ^5–9^ and have been visualised budding from intra-aortic clusters (IAHC) before being released into the circulation^10–14^. Research on the molecular mechanisms associated with the development of hematopoietic stem and progenitor cells (HSPCs) in the embryo has instructed the design of culture protocols to model hematopoiesis *in vitro* from pluripotent stem cells (PSCs)^15^.

However, it has proven very challenging to generate *bone fide* HSCs that are functionally capable of long term reconstitution and the small number of the reports that claim to have succeeded have only done so, albeit with low efficiency, by employing transgenic strategies and/or providing an in vivo environment for their maturation ^16,17, 18^. There is a great need to understand the cellular and molecular mechanisms associated with the development and maintenance of HSCs and to compare the key signaling pathways between *in vivo* and *in vitro* model systems.

The Apelin receptor gene (*Aplnr/Agtrl1/Apj*) encodes a member of the G protein-coupled receptor family and has been implicated in cardiac, endothelial and hematopoietic development in a number of model systems ^19–22^. Pertinent to this study is the involvement of Aplnr in endothelial cell maturation during development and the fact that it appears to be expressed at a higher level in endothelial cells of the AGM region, implying that it could be related to their hemogenic potential ^23 24, 25^. In the adult bone marrow endothelial cells that express Apelin, one of the Aplnr ligands, participate in the remodeling of the vascular following irradiation and drive post-transplant recovery through feedback from HSPCs^26^. The expression profile of Aplnr and its ligands Apelin and Apela during mouse embryonic development indicates that this signaling pathway is also active in mesoderm and its derivatives ^22, 27^. This is supported by findings in differentiating human ESCs that *APLNR* is expressed in Mixl1-expressing mesodermal progenitors and in transient mesenchymoangioblast cells, the common precursor to endothelial and mesenchymal cells^28 29^. Addition of Apelin to differentiating human PSCs increased the production of blast colonies that are considered to represent a common precursor to endothelial and hematopoietic cells ^28^. Our previous research has also implicated the Aplnr pathway during hematopoietic development in vitro. We demonstrated that expression of Aplnr and one of its ligands, Apelin, correlated with the increased production of HPCs when the transcription factor, HOXB4 was activated in differentiating mouse and human PSCs^30, 31^. Although all of these studies implicate a role for Aplnr signaling during hematopoietic development the specific function of this pathway has not been addressed directly. To do this, we generated Aplnr-null and Aplnr-dtTomato reporter murine embryonic stem cell (mESC) lines. We show here that the production of HPCs was significantly impaired in differentiating Aplnr-null ESCs and that the Aplnr-tdTomato reporter marked a population of differentiating mesoderm cells committed to hematopoietic and endothelial lineages. To assess the role of Aplnr signalling during the development of HSCs and hematopoietic progenitors and HSCs we added Apelin ligands to differentiating ESCs and to AGM explant cultures. No significant effect was observed on the production of HPCs from either mouse of human ESCs in vitro. However, in AGM explant cultures we noted a marked decrease in the number of functional LSK-HSCs and a concomitant increase in the production of myeloid cells.

## RESULTS

### Aplnr is required for hematopoietic cell production from ESCs

To assess whether Aplnr signaling was required during the differentiation of hematopoietic cells from mESC, we generated Aplnr null mESCs using a CRISPR/Cas9 strategy. *Aplnr* is a single exon transcript with a coding region of 1133 base pairs (bp) so our strategy involved excising the complete coding region using guide RNAs directed to the 5’ and 3’ ends (Supplementary Figure S1A). pSPCas9-2A-mCherry-U6-gRNA plasmids containing pre-selected gRNAs (see methods) were transfected into E14 mESCs. Cells that had been successfully transfected were sorted based on mCherry expression 24-48 hours after transfection then plated at low density. Individual mESC clones were isolated and genomic DNA was screened by PCR using primers that spanned the deleted region. The PCR assay was designed to amplify a 589 bp product from the deleted, knockout (KO) allele and a 1700 bp amplicon from the wild type (WT) allele. The KO allele was identified in 7 out of 30 mCherry-positive clones (Supplementary Figure S1B). This assay did not distinguish between heterozygous and homozygous clones because the smaller 589bp amplicon associated with the KO allele would likely be amplified preferentially over the larger WT allele. To differentiate between functionally heterozygous and homozygous mESC clones, we sequenced the amplicons and found that in the majority of clones that had one deleted *Aplnr* allele, the second *Aplnr* allele had an insertion or a deletion that resulted in a frame shift mutation (data not shown). Quantitative Real-Time PCR was used to select clones in which the targeting events resulted in ablation of *Aplnr* transcripts and could therefore be considered as functionally null at the *Aplnr* locus (Supplementary Figure S1C).

Aplnr-null mESCs could be maintained as undifferentiated ESCs in the presence of LIF in a comparable manner to controls (data not shown). However, when induced to differentiate and assessed in hematopoietic colony assays, significantly lower numbers of hematopoietic CFU-C colonies (CFU-M, CFU-GM and CFU-Mix) were formed from two independently derived *Aplnr*-null ESCs compared to control wild type ESCs at day 6 of differentiation (Figure 1A). Flow cytometry analyses confirmed that the Aplnr-null ESCs produced a lower number of CD41^+^VE-Cadherin^+^ HPCs (Figure 1B, C). When subjected to a macrophage-specific differentiation protocol, the proportion of mature F4/80^+^CD11b^+^CD16^+^ macrophages generated from *Aplnr*-null ESCs was significantly lower than controls (Figure 1D). To confirm that the reduced hematopoietic differentiation was due to the absence of Aplnr and not simply a consequence of ESC clonal variation, we introduced an Aplnr-expressing plasmid into the Aplnr-null ESC clones and demonstrated that the production of F4/80^+^CD11b^+^ macrophages was increased in the presence of the plasmid (Figure 1D). These data indicate that the Aplnr pathway is required for the development of HPCs and for the differentiation of the myeloid lineage.

**Figure 1.**
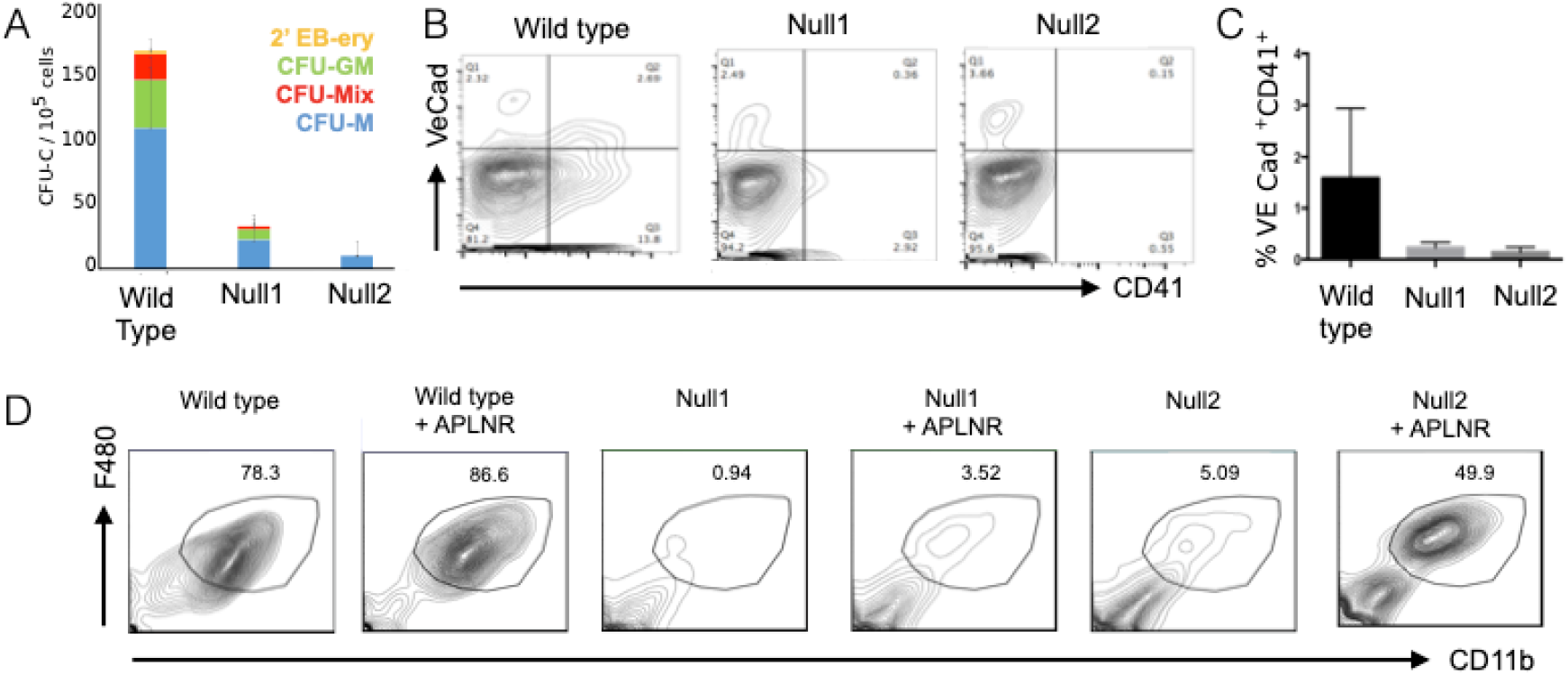
Targeted deletion of *Aplnr* results in impaired hematopoietic differentiation. A. The number of hematopoietic CFU-C colonies generated from 10^5^ differentiated (day 6) control ESCs and two independently-derived Aplnr-null (Null1 and Null2) ESC clones. CFU-M (macrophage), CFU-GM (granulocye/macrophage), CFU-GEMM (multi-lineage including granulocyte, erythroid, macrophage and megakaryocytes), 2’EB-ery (secondary embryoid bodies with associated erythroid cells). (n = 4 * p<0.05) B. Representative flow cytometry analyses of day 6 differentiated control ESCs and two independently-derived Aplnr-null ESC clones (Null1 and Null2) using antibodies to VE-Cadherin and CD41. C. Quantification of flow cytometry analyses (B) showing the percentage of VE-Cad^+^CD41^+^ cells at day 6 of differentiation from control and 2 Aplnr-null ESC clones (n= 3; * p <0.05) D. Representative flow cytometry of cells generated following the macrophage differentiation protocol of control (Wild type), two Aplnr-null ESC lines (Null 1 and Null 2) and the Null 1 and Null2 ESCs that were transfected with an Aplnr-expressing plasmid (+APLNR).

### Aplnr-tdTomato reporter ESC line tracks a transient mesoderm cell population

To define the phenotype of cells expressing the Aplnr receptor during hematopioetic differentiation of ESCs we generated an *Aplnr*-tdTomato reporter mESC line using CRISPR/Cas9-mediated genome editing (Figure 2A). mESCs were transfected with the *Aplnr* targeting vector, CAS9 plasmid and specific gRNAs. 35 G418 resistant colonies were selected and 18 of these were screened by genomic PCR and Southern blotting (Supplementary Figure S1D). We confirmed that the *Aplnr*-tdTomato reporter faithfully mimicked *Aplnr* transcript expression by demonstrating the expression of *Aplnr* transcripts in FAC-sorted tdTomato positive, but not tdTomato negative, cells (Figure 2B). However, we noted that the expression of the Aplnr-tdTomato reporter did not correlate with the expression of the Aplnr receptor that was supposedly detected by a commercially available α-APLNR antibody (Supplementary Figure S2). Real-Time PCR analyses of FAC-sorted cells detected *Aplnr* transcripts in cells sorted based on expression of the Aplnr-tdTomato reporter but not α-Aplnr antibody staining (Supplementary Figure S2B, C). This suggested that the α-Aplnr antibody was binding non-specifically to the cell surface and this was further confirmed by demonstrating that the α-Aplnr antibody detected a protein in our differentiating Aplnr-null ESCs (Supplementary Figure S2A) and in 293T that do not express Aplnr transcripts (Supplementary Figure S2D). These findings emphasized the requirement of the Aplr-tdTomato reporter mESC lines in defining the potential of Aplnr-expressing cells.

**Figure 2.**
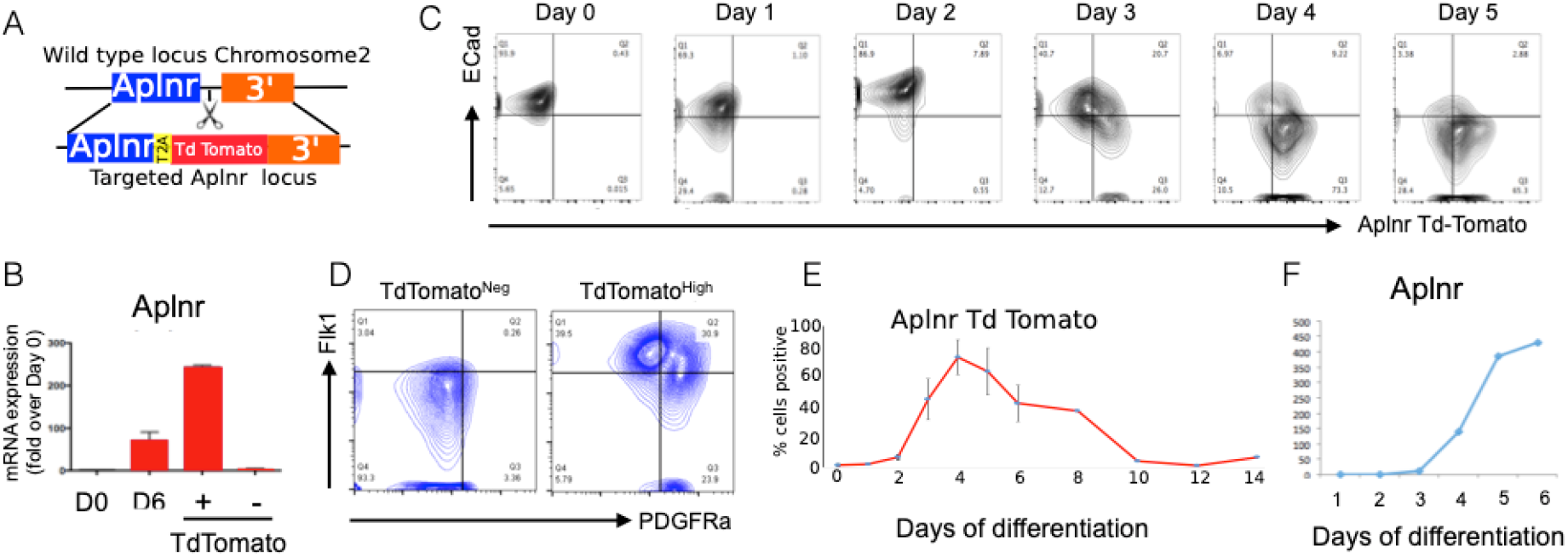
Aplnr-trTomato reporter ESC line mimics the activity of the Aplnr locus during ESC differentiation and tracks a transient mesoderm cell population. A. CRISPR/ CAs9 gene editing strategy. Guide RNAs were designed to cut the genomic region of chromosome 2 immediately after the *Aplnr* coding sequence. Schematic of targeting vector that was used to insert the tdTomato reporter gene at this site, followed by a T2A sequence and Neo^R^ cassette (not shown). B. Quantitative RT-PCR of *Aplnr* expression in undifferentiated Aplnr-tdTomato reporter ESCs (d0), unsorted differentiated day 6 (d6) cells and day 6 cells sorted into Aplnr-tdTomato positive (+) and negative (−) populations. (n= 3) C. Representative flow cytometry plots of Aplnr-tdTomato and ECadherin expression during differentiation from undifferentiated ESCs (day0 (d0) to day 5 of differentiation. D. Flow cytometry of Aplnr-tdTomato ESCs at day 3 of differentiation stained with antibodies to FLK1 and PDGFRα then analysed following gating on either APLNR negative or APLNR high. E. Percentage of Aplnr-tdTomato positive cells detected by flow cytometry in undifferentiated (day 0) cells and during a 14 day time course of differentiation. Error bars represent standard deviation of data from 3 independent experiments. F. Relative expression of *Aplnr* transcripts in differentiating ESC from day 0 to day 6 assessed by qRT-PCR.

The timing of expression of the Aplnr-tdTomato reporter during mESC differentiation was compared to the cell surface expression of E Cadherin (ECad), an epithelial marker that is expressed at a high level in undifferentiated pluripotent stem cells but downregulated upon differentiation into mesoderm lineages^32^. Since the APLNR is first expressed in the developing mesoderm of the mouse embryo, we would predict a mutually exclusive expression pattern when co-stained with ECad. Thus, as expected, we observed ECad expression in undifferentiated ESCs (day 0) and expression was gradually reduced as differentiation progressed (Figure 2C). In contrast, the *Aplnr*-Td-tomato reporter was not expressed in undifferentiated cells and expression increased throughout differentiation. *Aplnr*-tdTomato-expressing cells were first detected at day 3 of differentiation in both ECad^+^ and ECad^−^ cells but by day 5 virtually all the *Aplnr*-tdTomato-expressing cells were negative for ECad (Figure 2C). Thus, differentiating ESCs transition from ECad^+^Aplnr^−^ to Ecad^+^Aplnr^+^ phenotype then they lose expression of ECad to become Ecad^−^Aplnr^+^. The highest proportion (20%) of the intermediate, double positive cells was present around day 3. As further confirmation that the *Aplnr*-tdTomato reporter is expressed in the emerging multipotent mesoderm, we show that most of the cells expressing a high level of Aplnr-tdTomato at day 3 co-expressed high levels of Flk1 and around half of these also expressing PDGFRα indicating that this population was marking mesodermal cells (Figure 2D). The highest proportion (around 70%) of *Aplnr*-tdTomato cells was observed at day 4 and this gradually declined as differentiated proceeded (Figure 2E). The expression profile of *Aplnr*-tdTomato reporter expression in differentiating mESCs mimicked the expression of *Aplnr* transcripts in WT control cells that was virtually undetected in undifferentiated mESCs but increased during the course of differentiation (Figure 2F).

### Hematopoietic activity is within the cell population that express high levels of APLNR

*Aplnr*-tdTomato mESCs were differentiated into embryoid bodies (EBs) using a serum free hematopoietic protocol and at day 5 EBs were dissociated and cells expressing different levels of *Aplnr*-tdTomato were isolated by flow cytometry (Supplementary Figure S3). The differentiation potential of FAC-sorted cells was assessed using CFU-C assays (Figure 3A). Hematopoietic activity was found exclusively within in the *Aplnr*-tdTomato^+^ cell population with a significantly higher number of CFU-Cs being detected in cells expressing high levels of *Aplnr*-tdTomato compared to those expressing low levels. The majority of multilineage colonies (CFU-Mix) was associated with the cell population expressing the highest level of *Aplnr*-tdTomato (Figure 3A). Cells expressing the highest level of *Aplnr*-tdTomato also expressed high levels of the endothelial markers, VE Cadherin, Flk1, Tie2 and a subset of those also expressed the hematopoietic markers, CD41 and Kit, a cell surface phenotype associated with hemogenic endothelium (Figure 3B, C).

**Figure 3.**
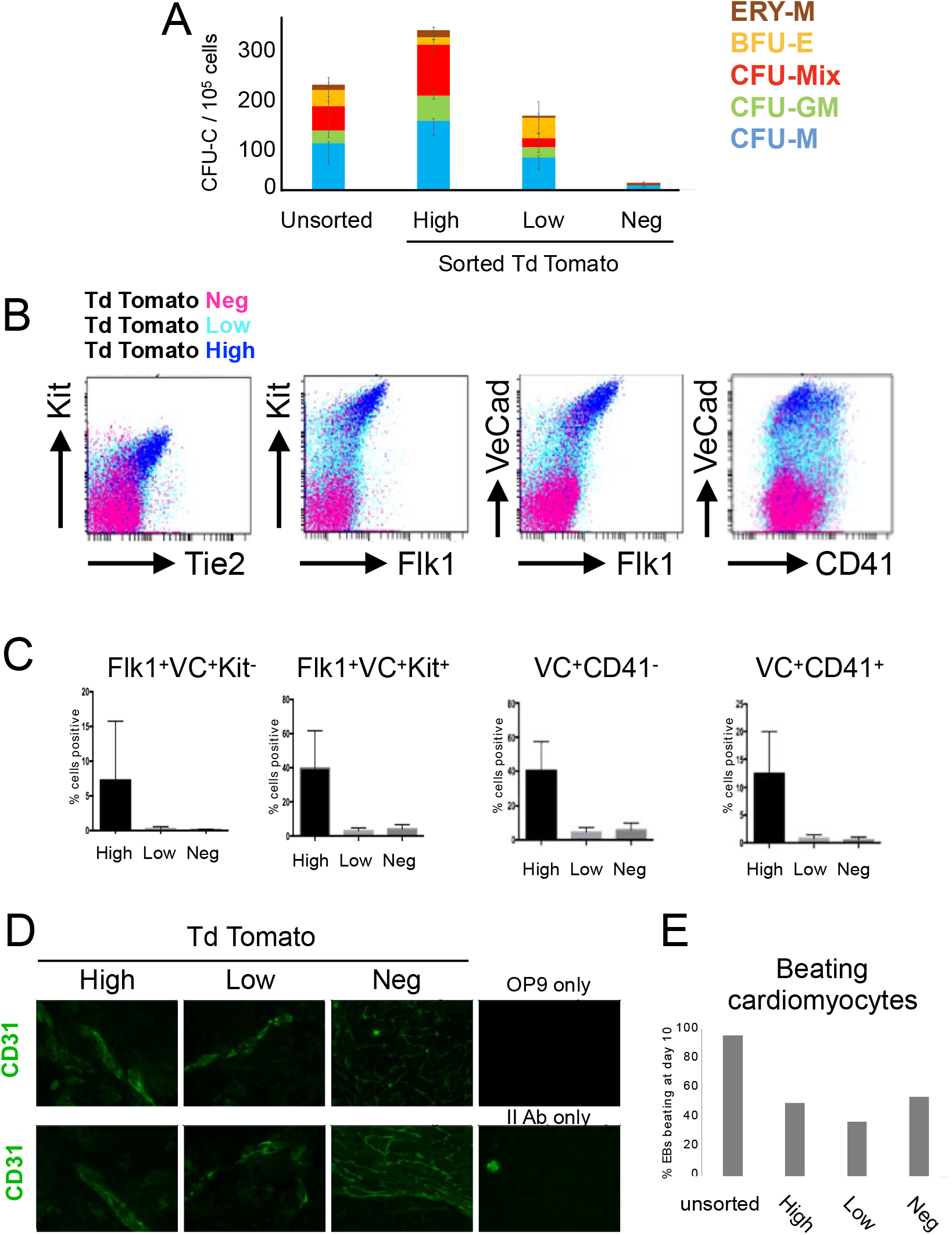
Aplnr-tdTomato marks mesoderm fated to become hemogenic endothelium. A. Number of CFU-Cs generated from 10^5^ unsorted differentiating ESCs (day 6) and cell populations sorted based a high (Hi), low (Lo) or negligible (Neg) level of tdTomato expression (See Supplementary Figure S3 for FAC sorting strategy) (n=3 * p<0.05)). B. Overlaid representative flow cytometry plots of differentiating Aplnr-tdTomato ESCs (day 6) co-stained with antibodies against Kit, Tie2, VE-Cadherin, Flk1 and CD41. Backgating of cells expressing high (dark blue), low (light blue) or negligible (pink) levels of Aplnr-tdTomato demonstrates that cells expressing Tie2, Flk1, VE-Cadherin (VC), Kit and /CD41 are primarily found in the Aplnr-tdTomato high cell population. C. Quantification of flow cytometry data in B showing the percentage of cell type within each of the sub-populations defined by the level of Aplnr-td-tomato expression (n= 3:*p<0.05)). D. Immunohistochemistry using an αCD31 antibody of differentiating ESCs following FAC-sorting and culturing on OP9 stromal cells in the presence of VEGF demonstrating the endothelial potential differentiating cells expressing high and low levels of Aplnr-tdTomato. Two replicate experiments are shown. E. Percentage of embryoid bodies (EBs) with associated beating cardiomyocytes in EBs generated from unsorted day 6 differentiating ESCs or cells at this stage that were sorted based on the expression of high (Hi), low (Lo) or negligible (Neg) levels of Aplnr-tdTomato expression.

To assess endothelial potential, *Aplnr*-tdTomato-expressing cells they were plated onto irradiated OP9 cells, cultured in the presence of VEGF for 10 days then immune-stained for the endothelial marker, CD31 (Figure 3D). Cells expression high and low levels of *Aplnr*-tdTomato generated endothelial structures where CD31^+^ cells were observed into vessel-like structure. No endothelial structures were produced by *Aplnr*-tdTomato negative cells. To assess whether the *Aplnr*-tdTomato reporter was marking all mesoderm progenitors or a subpopulation committed to hematopoietic and endothelial lineages, we assessed the potential of sorted cells to differentiated into another mesodermal cell type. EBs were dissociated at day 6, sorted based on the expression of *Aplnr*-tdTomato then assessed for their potential to generate cells of the cardiac lineage. Although sorting reduced the overall production of beating cardiomyocytes compared to unsorted cells, there was no correlation between the level of *Aplnr*-tdTomato expression and the potential to generate cardiac cells (Figure 3E). Taken together these data support our hypothesis that the *Aplnr*-tdTomato reporter is marking a specific subpopulation of hemato-endothelial biased mesodermal progenitors.

### Activation of the Aplnr pathway has no significant effect on the production of haematopoietic progenitors in differentiating mouse and human PSCs

Given that *Apnlr* is expressed on mESC-derived mesoderm committed to a hematopoietic and endothelial fate, we considered that activation of the Aplnr signaling by the addition of Apelin ligands might modulate the patterning of mesoderm and thereby alter the production of hematopoietic progenitors. When Apelin was added to the serum-free cytokine-based hematopoietic differentiation protocol we observed no effect on the number or phenotype of CFU-Cs that were generated (Figure 4A). This was somewhat unexpected given previous studies had reported to promote hematopoiesis when Apelin was added to differentiating human ESC^28^. To confirm that the effects we had observed were not species specific we replicated our findings using both human ESC and induced pluripotent stem cells (iPSCs) and again demonstrated that the addition of Apelin peptides had no effect on CFU-C formation (Figure 4B).

**Figure 4.**
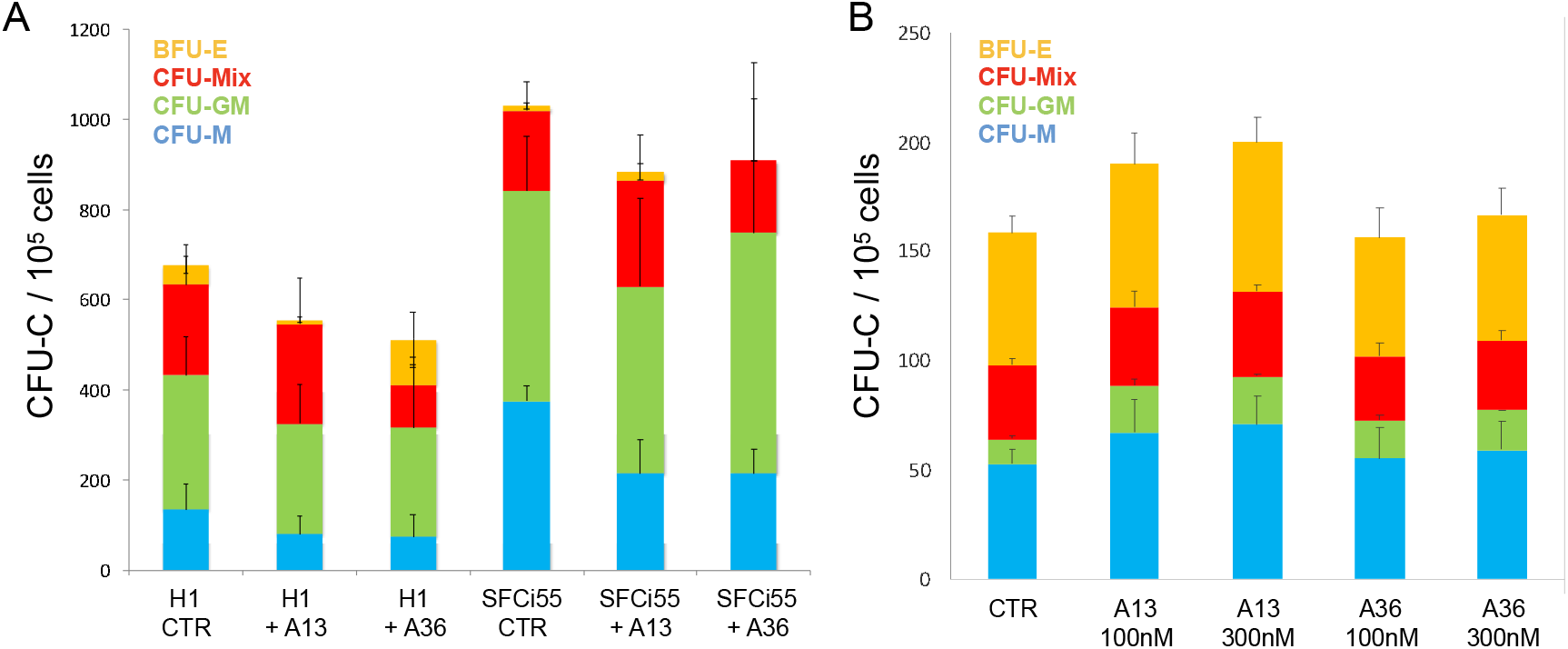
Addition of Apelin ligands to differentiating mouse and human PSC does not enhance hematopoietic colony production. A. Number of CFU-C generated in differentiating mouse ESCs with no Apelin ligands added (control) and following the addition of Apelin 13 (A13) or Apelin 36 (A36) at 30nM, 100nM (n=3) B. Number of CFU-C generated in differentiating human ES (H1) (n=8) and human iPSCs (SFCi55) (n=3) in control cultures with no Apelin ligands added (no A) and following the addition of Apelin 13 (A13) or Apelin 36 (A36).

### Aplnr and its ligands are expressed at the site of HSC emergence in vivo

As it is widely acknowledged that definitive HSCs are not efficiently generated in differentiating PSCs and the complex in vivo environment cannot be efficiently recapitulated in vitro, we studied the role of Aplnr in this complex process. To assess whether the Aplnr pathway could be active during the production of definitive HSCs in vivo, we first assessed the expression level of genes encoding the receptor and ligands in sites of hematopoiesis in the developing mouse embryo. *Aplnr* and genes encoding its ligands*, Apelin* and *Apela* were all expressed in the aorta-gonad-mesonephros (AGM) region of the developing embryo at day 9.5 and 11.5 when definitive HSCs develop and first emerge (Figure 5A). Low, but detectable levels of expression were also observed in yolk sac and foetal liver. These data support the hypothesis that the Aplnr signaling pathway is active during the emergence of definitive HSCs *in vivo*.

**Figure 5.**
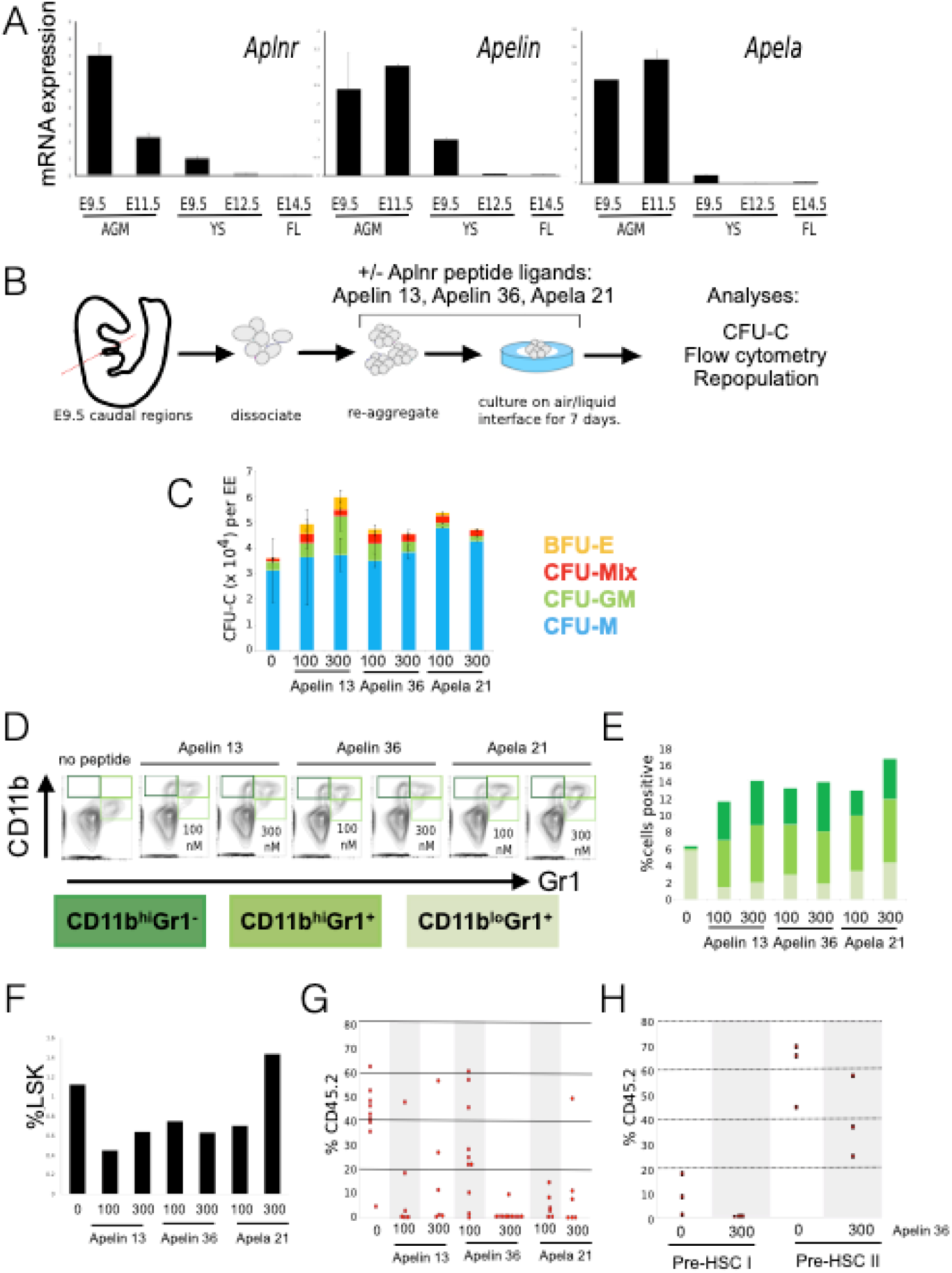
Addition of Aplnr ligands to aorta-gonad-mesonephros reaggregate explant cultures results in a reduction in transplantable HSCs. **A**. Quantitative RT-PCR analyses of RNA isolated from the aorta-mesonephros (AGM) region, yolk sac (YS) and foetal liver (FL) dissected from embryos at the indicated stage of development using primers to detect transcripts encoding Aplnr and its ligands, Apelin and Apela (n=2). **B**. Schematic of experimental strategy for E9.5 dissection (as example) where the caudal region of the embryo was dissected, tissue dissociated and set up in reaggregate cultures in the presence of Apelin peptides for 7 days then assayed by for colony formation (CFU-C), flow cytometry and transplantation into lethally-irradiated mice. **C**. CFU-C analyses of reaggregate cultures in the absence (0) or presence (100nM or 300nM) of Apelin 13, Apelin 36 or Apela 21 **D**. Representative flow cytometry plots of live cells obtained after 7 days of E9.5 aggregate culture in presence of Apelin 13, Apelin 36 or Apela 21 using antibodies against the myeloid markers, CD11b and Gr1). **E**. Quantification of flow cytometry in C. **F**. Proportion of lin^−^Sca^+^Kit^+^ hematopoietic progenitor cells after 7 days of E9.5 aggregate culture in the presence of Apelin 13, Apelin 36 or Apela 21 peptides. **G**. Proportion of CD45.2 donor cells present in the peripheral blood of lethally irradiated recipients 14 weeks after transplantation of cells derived from E9 ex vivo reaggregate cultures in the absence (0) or presence of 100 or 300nM Apelin 13, Apelin 36 or Apela 21. Each data point represents one recipient animal. **H**. Proportion of CD45.2 donor cells present in the peripheral blood of lethally irradiated recipients 14 weeks after transplantation of aggregate cultures that consisted of sorted pre-HSC-Type I and Type II cells derived from E11 in co-aggregate cultures in the absence (0) or presence of 100 or 300nM Apelin 13, Apelin 36.

### Activation of the Aplnr pathway reduces the production of transplantable HSCs in aggregate cultures

As transcripts encoding Aplnr and its ligands are expressed in the AGM region of the developing embryo this might indicate that the Aplnr pathway could influence the production, maintenance or differentiation HSCs. To test this hypothesis, we added Aplnr ligands, Apelin 13 (A13), Apelin 36 (A36) or Apela 21 (A21) to an established AGM explant reaggregation culture system that allows the ex-vivo production and maturation of definitive HSCs from their precursors (Figure 5B)^11, 33^. The caudal region of day 9.5 embryos that includes AGM tissue was dissected and placed in the aggregation culture in the presence of cytokines with or without Apelin peptides for 7 days. Resultant cells were assessed for CFU-C formation, flow cytometry and their ability to reconstitute lethally irradiated recipients *in vivo*. The addition of either A13 or A36 peptides to the aggregate cultures resulted in a slight, but not significant, increase in the total number of hematopoietic progenitors as assessed by CFU-C production (Figure 5C). Flow cytometry analyses of cells derived from the CFU-C colonies demonstrated an increase in the proportion of mature myeloid cells expressing high levels of CD11b, a proportion of which also expressed Gr1 (Figure 5D, E). The increase in the production of mature myeloid cells in the presence of Aplnr ligands coincided with a decrease in the production of immature (lin^−^/Sca^+^/Kit^+^ (LSK) hematopoietic progenitor cells (Figure 5F).

To assess the effects of Aplnr ligands on the production of functional reconstituting HSCs in *ex vivo* cultures, day 9.5 AGM cells (CD45.2) were cultured for 7 days in the presence of various concentrations of A13, A36 or A21 then resultant cells were transplanted into irradiated CD45.1/2 recipients. Almost all (8/9) recipients that received control cell aggregates cultured without Apelin peptides demonstrated successfully HSC reconstitution 14 weeks after transplantation (>10% CD45^+^). In contrast, when a high concentration (300nM) of either A13, A36 or A21 was added to the aggregate culture, the number of successfully transplanted animals was reduced to 3/6, 1/10 and 2/6, respectively (Figure 5G). The effect of peptide addition was dose dependent with the addition of a lower concentration (100nM) of all peptides having a less profound detrimental effect on the ability of cultured cells to reconstitute the hematopoietic system of irradiated recipient mice (Figure 5G). To define the timing of this detrimental effect, we isolated pre-HSC Type 1 (VC^+^CD41^lo^CD43^+^CD45^−^) and pre-HSC Type 2 cells (VC^+^CD41^lo^CD43^+^CD45^+^) from day 11.5 AGM tissue and assessed the effect of A36 on their subsequent maturation into functional HSCs within aggregate cultures (Figure 5H). In control cultures, pre-HSC Type 1 could be matured *ex vivo* into reconstituting HSCs but in the presence of Apelin 36 resultant cells did not reconstitute irradiated mice. A comparable, but less profound effect of A36 was observed on pre-HSC Type 2 when again reconstitution was significantly reduced. These data demonstrate that activation of the Aplnr pathway by the addition of ligands is detrimental to the generation or maintenance of functional HSCs in ex vivo cultures.

Taken together the results of our study indicate that the Aplnr pathway is required for the production of cells fated to become endothelial and hematopoietic lineages but the activation of the Aplnr pathway at a later stage negatively affects HSC production or maintenance. The increased production of myeloid cells in the presence of *Aplnr* ligands suggests that this signaling pathway favours differentiation over self-renewal.

## DISCUSSION

We previously identified the genes encoding Aplnr and its ligand Apelin as the most highly upregulated genes when HOXB4 was activated in differentiating mESCs ^30^. As this increase in expression correlated with an enhanced hematopoietic differentiation, we hypothesised that the Aplnr signaling pathway could be associated with the production and/or maintenance of HSPCs. Here we show that complete deletion of the *Aplnr* gene in murine ESCs impairs their ability to differentiate into both hematopoietic and endothelial lineages indicating that Aplnr signaling is involved at a stage earlier than the commitment to these lineages, such as the formation or subsequent patterning of the mesoderm. This is consistent with a previous report where knockdown of *Aplnr* expression by shRNA in differentiating mouse ESCs resulted in delayed expression of the pan mesodermal marker, *Bry* and a reduction in endothelial marker genes, *Tie1, Tie2* and *Flk1*^34^. Our finding that the *Aplnr*-tdTomato reporter marks mesoderm committed to hematopoietic and endothelial lineages is also consistent with previous studies in differentiating human ESCs where *APLNR* expression was enriched in Mixl1-expressing mesodermal progenitors and in transient mesenchymoangioblast cells, the common precursor to endothelial and mesenchymal cells ^28 29^. We show that the cells expressing the highest level of the *Aplnr*-tdTomato reporter also express the highest level of endothelial and hemogenic endothelial markers which is in keeping with a previous study in hESC where APLNR-positive cells, identified by binding of the fluoresceinated peptide ligand, were found in mesoderm populations and were enriched in hemangioblast colony-forming cells^28^. It is interesting to note in this context that a recent single cell sequencing study reported *Aplnr* expression in CD44^−^ endothelial cells but not in the slightly later, CD44^low^ Kit^+^ cell population that are considered to represent cells in transition towards a hematopoietic fate ^35^.

A role of Aplnr in the production of endothelial cells is indicated by our finding that endothelial structures can be generated from *Aplnr*-tdTomato-positive but not *Aplnr*-tdTomato-negative differentiating ESCs and by reports that Aplnr^−/−^ mutant mice die *in utero* and display numerous cardiovascular abnormalities including deformed yolk sac vasculature sac^25^. The early mortality of Aplnr^−/−^ mutant mice precluded a study of Aplnr signalling during hematopoietic development in vivo and this awaits a conditional knockout approach.

Our observation that different levels Aplnr signaling are likely required at the different stages of hematopoietic development and maintenance likely explains the apparent discrepancy between reports on the effects of Apelin ligands on the hematopoietic commitment of PSC. Addition of Apelin ligand to the early stages of human ESC differentiation increased the production of blast colonies ^28^ whereas we observed no change in CFU-C potential following the addition of Apelin ligands to mouse or human pluripotent stem cells. Interestingly it was reported that Aplnr was not expressed in CD34+ cells derived from umbilical cord blood that the addition of Apelin had no effect on the number or phenotype of colonies in CFU-C assays^28^.

In AGM explant reaggregation cultures we show that addition of Aplnr ligands results in a reduction in HSCs capable of long-term reconstitution and a concomitant increase in the production of more mature myeloid cells. This finding suggests that activation of the Aplnr pathway is detrimental to the maintenance of HSCs and provides the first indication that the precise modulation of Aplnr signalling is critical in steady state hematopoiesis. The increase in the production of myeloid cells by Apelin ligands in *ex vivo* cultures is consistent with our observation that the in vitro production of macrophages from ESCs is significantly impaired in the absence of Aplnr and that this deficiency can be rescued by transfection of an Aplnr-expressing plasmid.

The differential requirements for Aplnr signaling at precise stages of HSC emergence and maintenance is comparable to that reported for the Notch signaling pathway where stage-specific effects observed in production of adult HSCs in vivo ^36^ and in HSPC production in vitro from ESC ^37^.

Critical to the interpretation and the relevance of our findings to hematopoietic development and maintenance in vivo is the anatomical and temporal expression of Aplnr ligands. Pertinent to this point is the finding, using an *Aplnr-CreER Rosa26-mTmG* reporter mice, that Apelin is expressed in a sub-population of endothelial cells in adult bone. Conditional knockout of this endothelial cell population revealed their critical role in hematopoietic regeneration following myeloablation^26^. It is not clear whether which of the Apelin ligands are important in the activation of Aplnr in different cell types. A conditional knockout approach for the Aplnr receptor and the production of mice carrying the Aplnr-tdTomato reporter will be instrumental in gaining a fuller understanding the complexity of this pathway during hematopoietic development and maintenance in vivo.

## EXPERIMENTAL PROCEDURES

### Mouse ESC maintenance and differentiation

The mouse embryonic stem cell (ESC) line E14IV was maintained in Glasgow Minimum Essential Medium (Gibco) supplemented with 10% fetal calf serum (FCS) (Lonza), 2 mM Sodium pyruvate (Gibco), 4 mM L-glutamine (Gibco), 1% Non-essential amino acids (Gibco) and 0.1 mM β-mercaptoethanol (Sigma) and 100 U/ml of leukaemia inhibitory factor (LIF) as previously described ^30^.

### Hematopoietic differentiation of mouse ESCs

Two days prior to differentiation, mouse ESCs were plated in N2B27 medium supplemented with 100U/ml LIF and 10ng/ml BMP4 to allow cells to adjust to serum-free conditions. To initiate differentiation, at day 0 cells were disaggregated with Accutase (Gibco) then resuspended in N2B27 containing 2.5 ng/ml ActivinA, 10ng/ml bFGF and 5 ng/ml BMP4 and plated in ultra-low attachment plates (Stem Cell Technologies). Two days later, EBs were plated on gelatinized tissue culture plates in N2B27 containing 10ng/ml bFGF, 5ng/m BMP4 and 15ng/ml VEGF for 6 days, with a media change on day 4. At day 6 the cells were harvested using Accutase (Gibco), counted and single cell suspensions were stained with antibodies for flow cytometry and/or plated in CFU-C assays that were scored 10 days later. In experiments to assess the effects of activating the Aplnr pathway, Apelin peptides were added from day 0, specifically, Apelin 36 (LVQPRGSRNGPGPWQGGRRKFRRQRPRLSHKGPMPF), its cleaved bioactive pyroglutamyl form, (Pyr1) Apelin 13 (QRPRLSHKGPMPF) and Apela 21 (LYRHACPRRRCIPLHSRVPFP)(Phoenix Pharmaceuticals). Macrophages differentiation was carried out as described previous^38^.

### Human ESC and iPSC differentiation

hESCs and hiPSCs were maintained on CELLstart (Invitrogen) coated 6-well tissue culture plates (Costar) in StemPro medium comprising of DMEM/F12 with Glutamax (Invitrogen), 1.8% BSA (Invitrogen), StemPro supplement (Invitrogen), 0.1mM β-Mercaptoethanol (Invitrogen) and 20ng/ml human basic FGF (Invitrogen) as previously described ^39, 40^. Cells were passaged mechanically when they reached about 80% confluency using EZ passage tool (Invitrogen). In experiments to assess the effects of activating the Aplnr pathway, Apelin peptides were added from day 0.

### Generation of Aplnr Td tomato reporter ESC lines

AplnrT2ATdTomato HDR donor vector was based on the backbone pBSK-2A-iCre-frtneofrt (a gift from Heiko Lickert) which was digested with Not1 and EcoR1 to accept two overlapping PCR fragments and a 1Kb 3’Arm by Gibson assembly (NEB). Aplnr (NM 011784.3) is a single exon gene and so the coding region of approximately 1kb acted as the 5’ homology arm. A PCR fragment consisting of the Aplnr coding region and the T2A-encoding sequence was generated by amplification of BAC BMQ407G118 using primers overlapping the backbone at the 5’ end and introducing T2A sequence at the 3’end. T2ATdTomato sequence was generated by PCR of pTdTomato (Clontech). The integrity of the vector was tested by sub-cloning the entire expression cassette into CAG IRES Puro vector and the Tdtomato signal confirmed in transient transfected ESCs by flow cytometry.

Guide RNAs (gRNA) directed to the 3’ end of the Aplnr sequence were predicted by the Zhang lab algorithm ((https://zlab.bio). gRNAs directed to the sequence 3’ to the Aplnr stop codon were identified and tested functionally using the SplitAx assay^41^. Briefly, a SplitAx vector consisting of sequence encoding the N terminal of GFP, Aplnr sequence that covered the region that the gRNAs were directed to and sequence encoding the C terminal of GFP that was out of frame with the N terminal sequence. Following transfection of this SplitAx vector and gRNAs into HEK293 cells, gRNAs were selected if they successfully cut the SplitAx vector and resulted in reconstitution of an in-frame GFP. Transfection of gRNA A2 (TGGGTCAGACCCGCTGCACC) binding to CCTGGTGCAGCGGGTCTGACCCA and gRNA B2 (GGAGAAAGTACAGCCATGCT) binding to AGCATGGCTGTACTTTCTCC resulted in a positive GFP signal in the SplitAx assay and were used for the subsequent gene targeting. Guides A2 and B2 were cloned into TOPO blunt after a synthesized U6 promoter (IDT) before sub-cloning into the guide vector, pGL3-U6-sgRNA EGFP^42^. 2 × 10^6^ Juo9 (subclone of E14) murine ESCs were transfected in suspension with 3μg AplnrT2ATdTomato HDR donor vector, 1μg gRNA-A2 plasmid, 1μgRNA-B2 plasmid and 1μg Cas9 Nickase(D10A)-GFP plasmid (Addgene #48140) or Hu CAS9-GFP (Addgene #44720) using Xfect Stem transfection reagent (Clontech) in 100μl Xfect Buffer and 2μl Xfect polymer for 4 hours. mESC media was added to a volume of 2 ml and plating out overnight in a gelatinised 6 well. Following overnight incubation, 40,000 GFP^+^ cells were FAC-sorted and plated in a gelatinised 10cm plate. G418 selection (400μg/ml) was added the next day and after 7 days 28 G418-resistant colonies from the Cas9 D10A transfections and 80 colonies from the wild type Cas9 transfections were generated. 35 colonies were picked and screened by PCR for correctly targeted 3’ and 5’ ends then confirmed by Southern blot.

### Generation of Aplnr Null ESC lines

gRNA sequences were identified (http://www.sanger.ac.uk/htgt/wge/find_crisprs). Four gRNAs with PAM sites after the ATG that had a low number of predicted off target binding sites and two guides with the PAM site after the TAA stop codon were selected. Linker nucleotides were added to the ends of the gRNA sequences and both strands synthesised (IDT). gRNAs were annealed and then ligated into the Bbs1 site of linearized pSPCAs9(Guide)-2A-mCherry vector. Combinations of 5’ and 3’ gRNAs were tested by transient transfection of mESCs and screening for excision of the coding region by PCR. Combinations with the highest level of excision in these transient transfections were used to generate clonal Aplnr null ESC. 1×10^6^ E14 ESCs were transfected with the two appropriate pSPCAs9(Guide)-2A-mCherry vectors using Xfect (Clontech). 48 hours after transfection cells were sorted for high levels of mCherry expression and plated onto 10 cm gelatinized dishes. Ten days later individual colonies were picked and screened using primers that spanned the coding region.

### Animals

Staged embryos were obtained by mating C57BL/6 (CD45.2/2) and the morning of discovery of the vaginal plug was designated as day 0.5. For culture and transplantation embryo E9.5 (25–29 sp), E11.0–E11.5 (>40 sp) were used. All experiments with animals were performed under Project License granted by the Home Office (UK), University of Edinburgh Ethical Review Committee, and conducted in accordance with local guidelines.

### Long-Term Repopulation Assay and blood chimerism analysis

CD45.2/2 cells were injected into irradiated 2-3 month old Bl/6J CD45.1/2 heterozygous recipients along with 100,000 CD45.1/1 nucleated bone marrow carrier cells per recipient. Recipients were □-irradiated using two doses (600 + 550 rad) separated by three hours. For day 9.5 culture one embryo equivalent (e.e.) of the 7 day cultured cells was injected. E11 AGM sorted cells where injected after 5 days OP9-coaggregate culture at a dose of 1e.e./recipient. Donor-derived chimerism was evaluated in peripheral blood at 6 and14 weeks post transplantation. Erythrocytes were lysed using PharM Lyse (BD Bioscience) and non-specific binding was blocked cells with an anti-CD16/32 (Fc-block) followed by staining with anti-CD45.1-APC (cloneA20) and anti-CD45.2-PE (clone 104) monoclonal antibodies (eBioscience). Cell populations were identified as CD45.1-PE+ (donor), CD45.1-PE+CD45.2-APC+ (recipient), CD45.1-APC+ (carrier) using FACSCalibur or Fortessa (BD Biosciences).

HSC numbers were assessed using ELDA analysis (Hu and Smyth, 2009). Multilineage donor-derived hematopoietic contribution in recipient blood and organs was determined by staining with anti-CD45.1-V450, anti-CD45.2-V500 and lineage-specific anti-Mac1-fluorescein isothiocyanate (FITC), Gr1-PE CD3e-APC, B220-PE-Cy7 monoclonal antibodies (BD Pharmingen).

### Ex vivo maturation of HSC precursors

E9.5 and E11.5 embryos were dissected as described^33^. Tissues were incubated with collagenase/dispase solution (0.12 mg/ml)(Roche) at 37° as described^11^, washed and resuspended in PBS (Sigma) containing 3% FCS then dissociated by pipetting. After dissociation (and sorting) 1ee of the specific cell populations (eg type 1 or type 11 pre-HSCs) were co-aggregated with 10^5^ OP9 cells. 5-10 co-aggregates (ie 5-10e.e.) per experimental variant were cultured in Iscove’s modified Dulbecco’s medium [IMDM], (Invitrogen), with 20% of preselected, heat-inactivated FCS, L-glutamine, penicillin/streptomycin) supplemented with murine recombinant cytokines (SCF, IL3 and Flt3) each at 100 ng/ml (PeproTech) and various concentrations of Aplnr ligand peptides including Apelin 36 (LVQPRGSRNGPGPWQGGRRKFRRQRPRLSHKGPMPF), its cleaved bioactive pyroglutamyl form, (Pyr1)Apelin 13 (QRPRLSHKGPMPF) and Apela 21 (LYRHACPRRRCIPLHSRVPFP)(Phoenix Pharmaceuticals). Co-aggregates were cultured on floating 0.8 μm AAWP 25 mm nitrocellulose membranes (Millipore) for 6 days then dissociated using collagenase/dispase as described ^33^.

### CFU-C and endothelial assays

Methylcellulose-based hematopoietic progenitor assay was routinely used to enumerate haematopoietic progenitors from both mouse (MethoCult™ GF M3434, Stemcell Technologies) and human (MethoCult™ H4434, Stemcell Technologies) differentiating PSC. Cells were collected from the differentiating PSC culture and prepared into single cell suspension. 1.5ml of methylcellulose medium were added into a 35mm low attachment dish. For each cell treatment two dishes were set up in parallel at densities 1×10^5^ and 5×10^4^ for mouse cells and at 1×10^4^ and 5×10^3^ for human cells then incubated at 37°C then scored between 7-11 or 12-14 days for mouse and human respectively. Colonies were classified based on the morphology using light microscopy. For endothelial assays, cell was placed on OP9-cell layer in presence of 50ng/ml VEGF as described ^33^. After 11 days, cultures were stained with anti-CD31 antibodies to assess of endothelial colonies.

### Fluorescence-Activated Cell Sorting and Analysis

Cell suspensions were stained with following antibodies: Ter119-v500 clone (Clone TER-119); anti-CD41-BV421 (brilliant violet 421) or Alexa Fluor 488 (clone MW30reg); anti-CD45 FITC (clone 30-F11); biotinylated anti-VE-cadherin (clone 11.D4.1) followed by incubation with Streptavidin-APC (all purchased from BD Pharmingen or Biolegend). Anti-mouse VE-cadherin (VC) antibody was biotinylated in-house using FluoReporter Mini-Biotin-XX Protein labeling kit (Invitrogen). Cell populations were sorted using a FACSAria-II sorter (BD) followed by purity checks. Type-I pre-HSC (Ter119^−^VC^+^CD41^+^CD45^−^) and type-2 pre-HSC (Ter119^−^VC^+^CD41^+^CD45^+^) were sorted from E11 AGMs as described previously ^7^. Dead cells were excluded using 7AAD staining and fluorescence minus one (FMO) control staining was used to define gating strategies. Data acquisition and analyses were performed using Fortessa (BD) and FlowJo software.

### Statistics

Data on histograms presented as average of at least three independent experiments ± SD and difference evaluated using t test or Prism software.

## AUTHORS CONTRIBUTIONS

MJ, AHT, RA, AF, SR and SM performed experiments. MJ and LF planned the experiments and wrote the manuscript. AM and TB provided intellectual input to the execution of the experiments and interpretation of the results

## ACKNOWLEDGEMENTS

This work was funded by the Wellcome Trust (AF, AHT) and Bloodwise (MJ). We thank Fiona Rossi and Claire Cryer for flow cytometry.

