## Supplementary Figures for "Modulation of Aplnr signaling is required during the development and maintenance of the hematopoietic system"

**Jackson et al Supplementary Figures**

Supplementary Figure S1

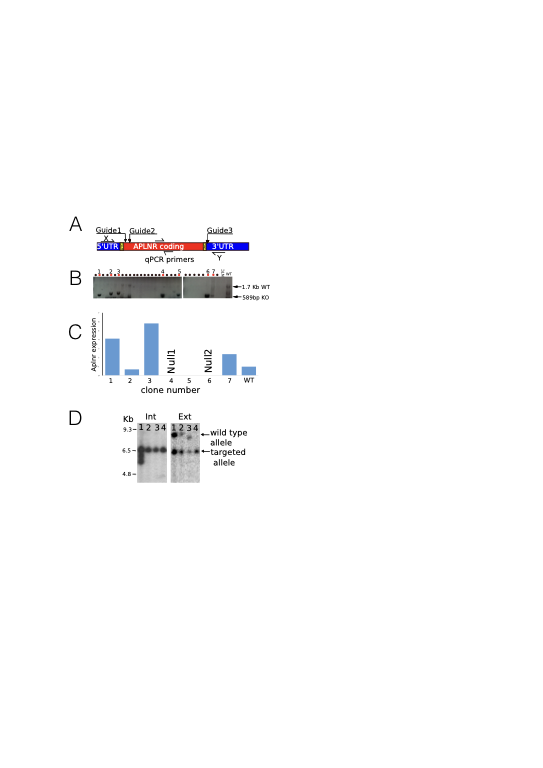

**Supplementary Figure S1**

A. Schematic of Crispr/Cas9 strategy to knockout the Aplnr coding region showing guide RNAs (Guide 1, Guide 2) and primers that were used to genotype resultant ESC clones and to validate depletion of Aplnr transcripts in qRT- PCR experiments.
B. Southern blot showing the presence of the wild type, 1700 base pair restriction fragment in control ESC and the predicted 589 base pair fragment in targeted clones.
C. qRT-PCR of the parental control ESC line (WT) and individual targeted ESC clones (1-7) showing that 2 of these (clones 4 and 6) had no APLNR transcripts, confirming functional homozygosity. These were then defined as Null 1 and Null 2, respectively.
D. Southern blotting screening of clones from Aplnr-tdTomato reporter ESC line production. Genomic DNA of G418-resistant clones was performed using a probe directly to the Neo sequence. Clones 2, 3 and 4 demonstrated a single integration of the targeting vector into the correct site.

Supplementary Figure S2

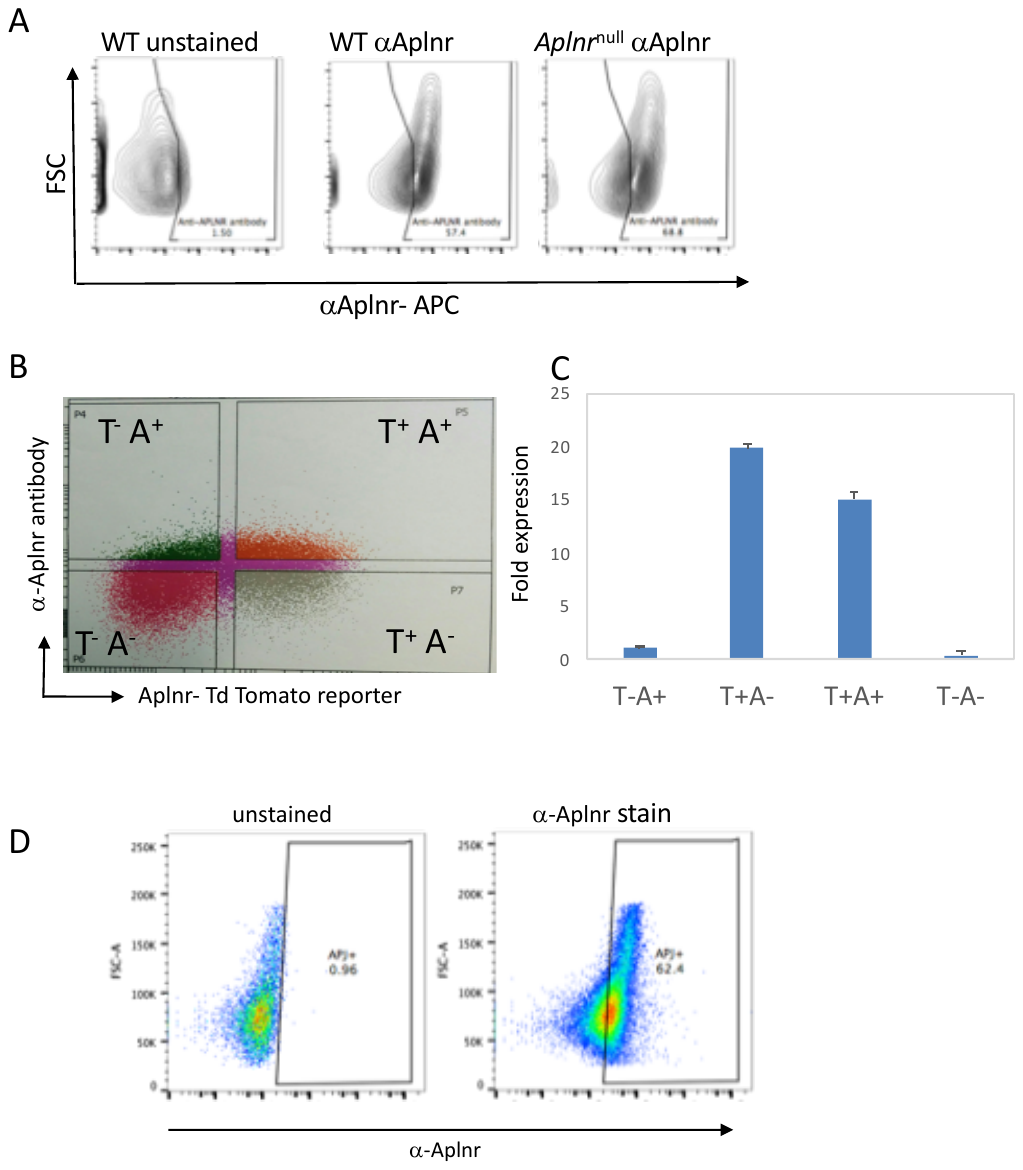

**Supplementary Figure S2. Commercial α-Aplnr antibody is not specific for Aplnr.**

A. Flow cytometry analyses of control, wild type (WT) ESCs and Aplnr-null ESCs stained with the anti-Aplnr antibody and unstained control.
B. Flow cytometry analyses of the Aplnr-tdTomato reporter ESC line stained with a commercial antibody to Aplnr.
C. qRT-PCR analyses of FAC-sorted cells based on the expression of the tdTomato reporter (T) and a-Aplnr antibody staining (A). The presence of *Aplnr* transcripts correlate with the tdTomato expression but not with cells identified by the Aplnr-antibody.
D. Control flow cytometry analyses of 293T cells stained with the Aplnr antibody demonstrates the poor specificity of the α-Aplnr antibody as the 293T cell line does not express Aplnr transcripts.

Supplementary Figure S3

**Supplementary Figure S3. Gating strategy for Aplnr-expressing cells**

Flow cytometry plot demonstrating the gating for sorting of Aplnr high , Aplnr mid and Aplnr negative cell populations. Insert show the re-analysis of sorting cells demonstrating purity of each of the populations.
